# What challenges remain in harmonizing cytomegalovirus viral load quantification across laboratories?

**DOI:** 10.1101/2024.12.09.627549

**Authors:** D. Boutolleau, A-S. L’Honneur, R. Germi, B. Chanzy, C. Archimbaud-Jallat, C. Rzadkowolski, J.B. Raimbourg, D. Gauthier, V. Thibault

**Affiliations:** AP-HP. Sorbonne Université, Pitié-Salpêtrière Hospital, Virology Department, National Reference Centre for Herpesviruses (Associated Laboratory), F-75013 Paris, France; Sorbonne University, INSERM, Pierre Louis Institute of Epidemiology and Public Health (IPLESP), UMR_S1136, F-75006 Paris, France 2Department of Virology, Cochin Hospital and University of Paris Cité, AP-HP, F-75014, Paris, France; Laboratoire de Virologie, CHU Grenoble, University Grenoble Alpes, CNRS, CEA, IRIG IBS, F-38000 Grenoble, France; CH Annecy Genevois, F-74000 Annecy, France; Service de Virologie CHU Clermont-Ferrand, Centre National de Référence des entérovirus et parechovirus, F-63000 Clermont-Ferrand, France; Bio-Rad Laboratories, Fort Worth, TX (USA); Marnes-La-Coquette (France); Irvine, CA, USA; CHU Rennes, Virology, F-35000 Rennes, France; Univ Rennes, Inserm, EHESP, Irset (Institut de recherche en santé, environnement et travail) UMR_S 1085, F-35000 Rennes, (France)

**Keywords:** calibration, Viral load monitoring, Extraction, Fragmentation, immunosuppression

## Abstract

Cytomegalovirus (CMV) infection monitoring is a key element in the management of immunocompromised patients. CMV DNA quantification in plasma or whole blood is the best indicator for clinicians to adjust immunosuppressive or antiviral therapies. Despite the availability of internationally standardized material, the commutability of CMV quantification results across laboratories remains inadequate. To assess inter-laboratory variability in CMV DNA quantification, we conducted a blinded study in seven independent laboratories. Each participant received a panel of 92 specimens for CMV quantification using their routinely used standard platform. While quantifications were highly correlated and reproducible, large discrepancies were observed with differences up to 1.45 log_10_ IU/mL between techniques for identical specimens. However, quantification scattering was lower for the WHO international standard or a commercially tested control (IQR=0.129) than for clinical specimens (0.469; p=0.0142). Blind quantification of the WHO or the commercial standard indicated that all techniques, except for fully integrated platforms, did not align well with the expected values and most platforms tended to quantify specimens and standards differently. Recalibration of all platforms against the same standard improved the spread of results, but differences of up to 1.19 log_10_ IU/mL remained for the same specimens. Achieving commutability in CMV quantification remains an elusive goal. Efforts should focus on improving both the assay calibrators and the run controls, which currently do not appear to simulate the unique characteristics of circulating CMV in patients. Until this is resolved, each transplanted patient should be consistently monitored by the same laboratory on the same platform.

## Introduction

Cytomegalovirus (CMV) is a herpesvirus (also known as human herpesvirus 5 or HHV-5) that contains the largest genome of any virus known to infect humans (1). CMV is very common worldwide in people of all ages. By the age of 40, more than half of the population is thought to be infected with this virus. CMV can spread through a variety of body fluids (saliva, respiratory secretions, urine, blood, tears, semen, genital secretions, and breast milk) as well as through blood transfusions and organ transplants from infected donors. After infection, CMV establishes latency for the life of the host and remains under the control of the immune system in healthy individuals, preventing it from causing disease or symptoms. However, in situations where the immune system is impaired, CMV can reactivate and lead to overt clinical manifestations (1). Transplant patients are at risk of developing allograft rejection which can sometimes be fatal. There is no sterilizing cure for CMV, but antiviral treatments help manage the infection when properly monitored by viral load quantification (2, 3).

Establishing the CMV serological status of donors and recipients is a prerequisite for transplantation in order to assess the need for and type of treatment strategy. In fact, the highest risk of CMV-associated end-organ disease arises when the recipient has never been exposed to CMV in his or her lifetime and has no pre-existing immunity. To minimize the risk of end-organ disease, it is also critical to monitor the recipient’s CMV viral load (VL) over time after transplantation to determine if preemptive antiviral therapy is needed and to apply the appropriate protocol (4).

Quantitative real-time PCR (qPCR) is the method of choice for monitoring CMV VL. This technique measures active CMV replication in the blood as latent CMV is typically undetectable by this method (5). Because qPCR quantifies the levels of nucleic acid present, whether from infectious or noninfectious virions, it measures DNAemia directly and viremia indirectly. Depending on the country and the laboratory, CMV DNAemia is measured in either whole blood or plasma (6). CMV DNA is present both intracellularly and cell-free in whole blood, whereas it is mostly cell-free and highly fragmented in plasma. Despite similar kinetics and peak infection times in both matrices, CMV DNAemia differs between them. In fact, CMV DNA levels are on average 1 log10 IU/mL higher in whole blood, especially during the ascending phase of infection. This elevated level allows for earlier detection of CMV DNA and therefore earlier initiation of antiviral treatment, although this increased sensitivity may sometimes lead to unnecessary treatment. Conversely, plasma CMV DNA levels are higher in the descending phase of infection, and this slower decay of CMV DNA may delay treatment interruption when antiviral therapy is administered (6).

Whether whole blood or plasma is chosen for testing, it is crucial to use the same matrix continuously to ensure consistent intra-laboratory results (7). The problem is that it is common for patients to be followed by more than one laboratory over time. In this case, although the same matrix may be tested, it is not sufficient to ensure that the measured CMV VLs are commutable between the laboratories monitoring these patients. Until 2010, all commercially available nucleic acid amplification tests (NAATs) reported CMV DNAemia in copies or genome equivalents (GE), which are units that are meaningful only within the context of a given assay but are not commutable across assays. To address this issue and help harmonize CMV results between laboratories, the World Health Organization (WHO) launched the 1st International Standard (IS) for CMV in 2010. This initiative was based on an international collaborative study that led to the creation of an International Unit (IU) that represents a “universal” consensus output resulting from the testing of the same CMV material by 32 different laboratories worldwide using a variety of routinely used NAATs (8). The WHO IS is considered the primary standard and is intended to be used by manufacturers and laboratories to calibrate their secondary standards. While this improves inter-laboratory commutability, it still does not allow for full harmonization, highlighting that the commutability issue is multifactorial and goes beyond the effects of matrix and reporting scale (9, 10). Other factors such as the calibration methodology (digital PCR vs qPCR) against the WHO primary standard, the type of secondary standard (whole virus *vs* purified genome or partial sequences) and matrix (clinical *vs* synthetic), the variability in extraction efficiencies, the gene target chosen, and the spacing between primer sequences all contribute to the lack of full harmonization (9).

Comparison of the in-kit standards of 13 commercial CMV qPCR assays revealed that none are made from whole virus, most are in a synthetic matrix, some are not calibrated to the WHO IS, and nearly half do not require extraction. Therefore, none of these CMV qPCR assays use standards that simulate patient specimens to encounter the same challenges and limitations. To assess the importance of using standards that closely mimic patient specimens, we ran the same CMV panel (including CMV clinical specimens, WHO IS, and Bio-Rad commercial standard and controls) along with in-kit controls in seven independent laboratories, each using its own and unique CE-IVD system (combination of assay and platform). We then compared the intra- and inter-laboratory results obtained (i) with the corresponding in-kit standards used as calibrators; (ii) after re-calibration with a Bio-Rad Exact Diagnostics (EDX) commercial CMV standard; and (iii) for the first time, after re-calibration with the current WHO IS.

## Material and methods

### Study design

Seven sites (six laboratories from France and one manufacturer from the US) were invited to participate in a multicenter CMV VL quantification study. Laboratories were selected based on their ability to measure CMV VL in IU/mL from plasma samples using a commercially available system. The selected laboratories reflect the heterogeneity of systems available in France. All laboratories were asked to blindly determine the CMV VLs from a panel of 92 samples, prepared as follows: 100 clinical plasma specimens, with previously quantified CMV VL, were sourced from a commercially-available collection (Inospecimens BioBank). Forty-three specimens randomly selected from this collection, along with the CMV WHO IS, were diluted to cover a broad concentration range as described below, and to ensure sufficient volume for inclusion in identical panels. These dilutions were made from a pool of CMV-negative plasma specimens, confirmed as negative by serology and PCR. Seven identical panels of 92 samples were then assembled that contained (i) 30 duplicates of CMV-positive clinical specimens (ranging from 2.0 to 6.0 log_10_ IU/mL); (ii) 3 duplicates of CMV-negative clinical specimens; (iii) 5 duplicates of the 1^st^ CMV WHO IS at 2.3, 3.6, 4.6, 5.6 and 6.6 log_10_ IU/mL (referred to as the WHO standard in this study); (iv) 5 duplicates from Bio-Rad’s Exact Diagnostics Verification Panel 1.4 mL (CMVP200) at the same concentration levels as for the WHO standard (referred to as the EDX standard); (v) 1 duplicate of the Bio-Rad’s Exact Diagnostics CMV Low Run Control (CMVL101) at 3.78 log_10_ IU/mL; (vi) 1 duplicate of the Bio-Rad’s Exact Diagnostics CMV High Run Control (CMVH102) at 5.7 log_10_ IU/mL; and (vii) 1 duplicate of PCR-grade water. All the panel members were stored at −20°C before shipment, then sent blinded to the participating laboratories.

### Commercial systems

The various systems used in this study are summarized in Supp. Table 1. Seven distinct workflows are represented, underscoring the various combinations of extraction methods with NAATs or integrated platforms.

### Statistical analyses

All results were initially generated using the manufacturers’ in-kit standards and analyzed in a blinded fashion. Blinding was then removed and re-quantification was performed sequentially using the WHO and the EDX standards. Since there is no gold standard method for CMV VL quantification, the results obtained on each system after each calibration were compared to the mean of all corresponding results for each of the panel members. The measured VL for the WHO and EDX standards were compared to their published nominal values. All participants reported their results in both IU/mL and Ct values. After blinding removal, the Ct values associated with the WHO and EDX standards were used to generate new standard curves for each system, and the VL of each panel member was calculated using these new calibration curves instead of the in-kit calibration.

Results are reported as median and 25th-75th percentile interquartile range (IQR), mean with standard deviation or numbers with percentages, as indicated. The Fisher exact test and chi-square test were used for categorical variables, where appropriate. The Mann-Whitney U test was used for continuous variables.

## Results

### Blind quantification of the WHO and EDX standards

Figure 1 shows the correlation between the nominal and measured values of the WHO standard when quantified by different systems. The nominal WHO standard VLs are 2.3, 3.6, 4.6, 5.6 and 6.6 log_10_ IU/mL. The measured values represent the mean of duplicates obtained with each system. The mean of the differences between the obtained and expected values ranged from −0.07 log_10_ IU/mL (from −0.42 to 0.06) for Alinity to 0.68 log_10_ IU/mL (from 0.57 to 0.76) for Ingenius (p<0.001). The slopes were generally below but close to 1 (from 0.91 to 1.01), indicating good linearity in the quantification of the WHO standard over its concentration range on the different systems. The same observation applies to the EDX standard (data not shown). Overall, the quantifications were highly correlated over the entire concentration range of both the WHO and the EDX standards.

**Figure 1:**
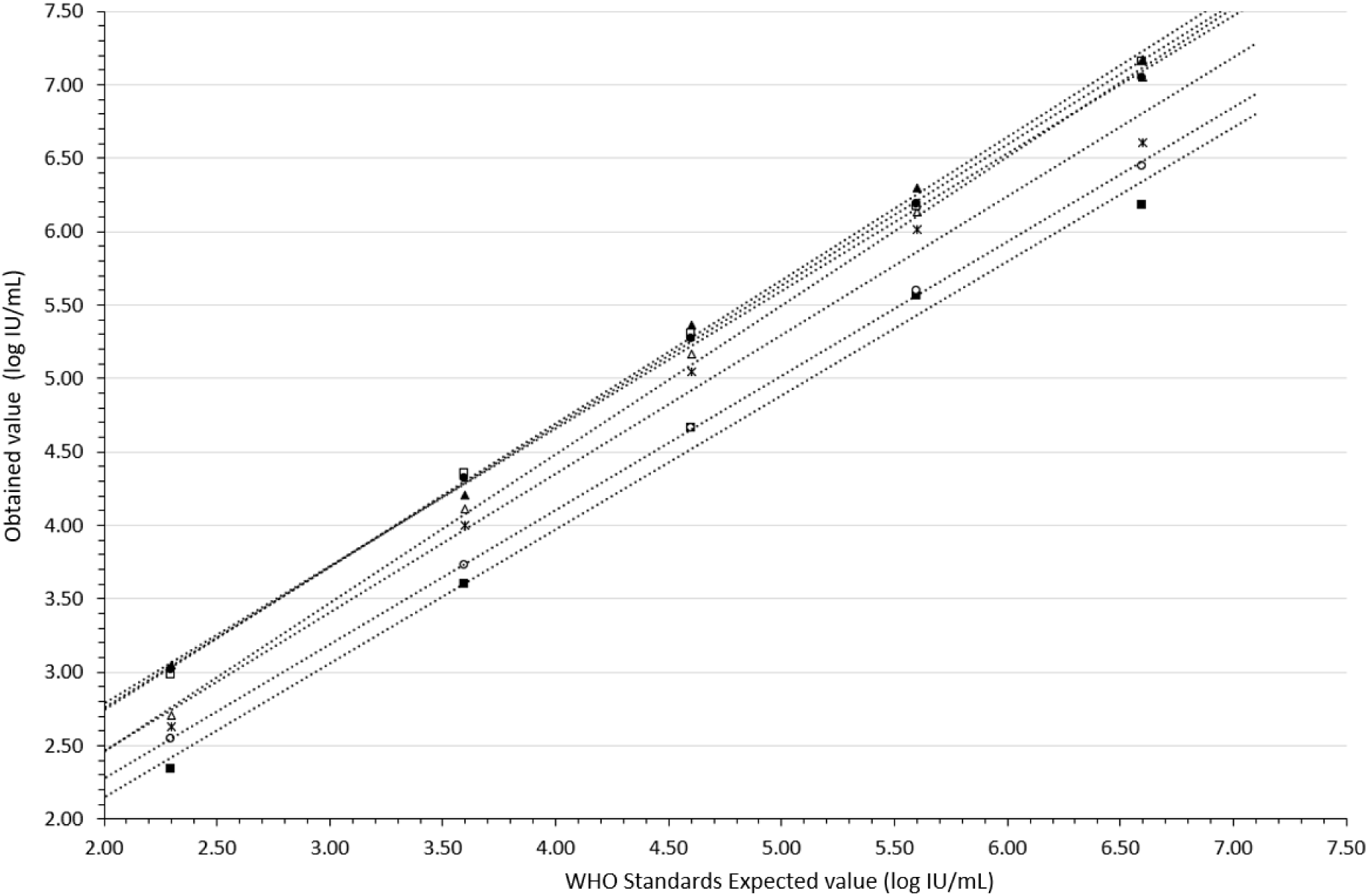
Correlation between the nominal and measured values of the WHO standard when quantified by different systems. Closed squares: Alinity; Closed triangles: Ingenius; Closed circles: Ingenius-Altona; Open triangles: Nuclisens-Rgene; Stars: Qiagen-Artus; Open circles: Roche; Open squares: Roche-Altona

### Blind quantification of the clinical specimens

The clinical specimens were blindly quantified with all systems, and because there is no gold standard for CMV VL quantification, the results were compared to the overall mean obtained of all systems combined. The quantification corresponded to the mean of the duplicates measured by each system. As observed for the WHO and EDX standards, the VLs were highly correlated between systems over the concentration range (Figure 2). Correlation slopes ranged from 0.82 for Roche to 1.06 for Ingenius. For each clinical specimen, the scattering of quantifications extended from 0.39 to 1.45 log_10_ IU/mL, across the concentration range.

**Figure 2:**
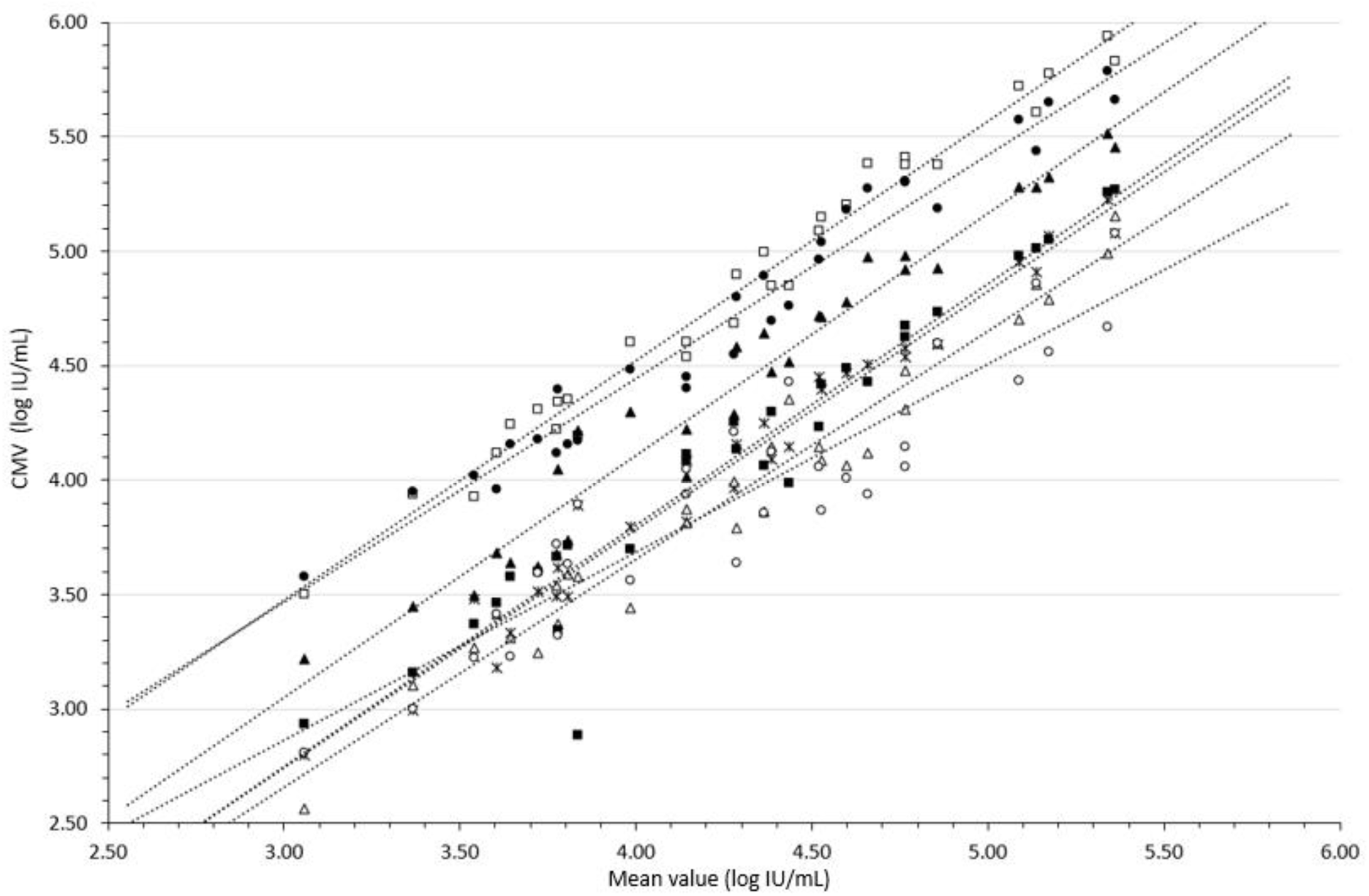
Distribution of VL obtained on clinical specimens. The x-axis represents the mean VL of each clinical specimen as measured by each system. The y-axis represents the overall mean VL obtained from all the systems combined. Closed squares: Alinity; Closed triangles: Ingenius; Closed circles: Ingenius-Altona; Open triangles: Nuclisens-Rgene; Stars: Qiagen-Artus; Open circles: Roche; Open squares: Roche-Altona

The differences between duplicate measurements of the clinical specimen VLs ranged from 0.06 +/-0.05 (SD) to 0.10 +/-0.11 log_10_ IU/mL, depending on the system (Figure 3). The maximum observed difference between two duplicate measurements once reached 0.75 log_10_ IU/mL (System Qiagen-Artus, duplicate sample with mean VL = 4.35 log_10_ IU/mL) but the overall mean difference between duplicates across all systems was quite narrow at 0.08 log_10_ IU/mL. Therefore, except for the aforementioned outlier, reproducibility did not differ significantly between systems, and an inverse correlation between reproducibility and VL was not systematically identified for each system. For the Roche-Altona system specifically, a trend (r=-0.626) was observed resulting from slightly greater differences between duplicates at the lower end of the quantification range (data not shown).

**Figure 3:**
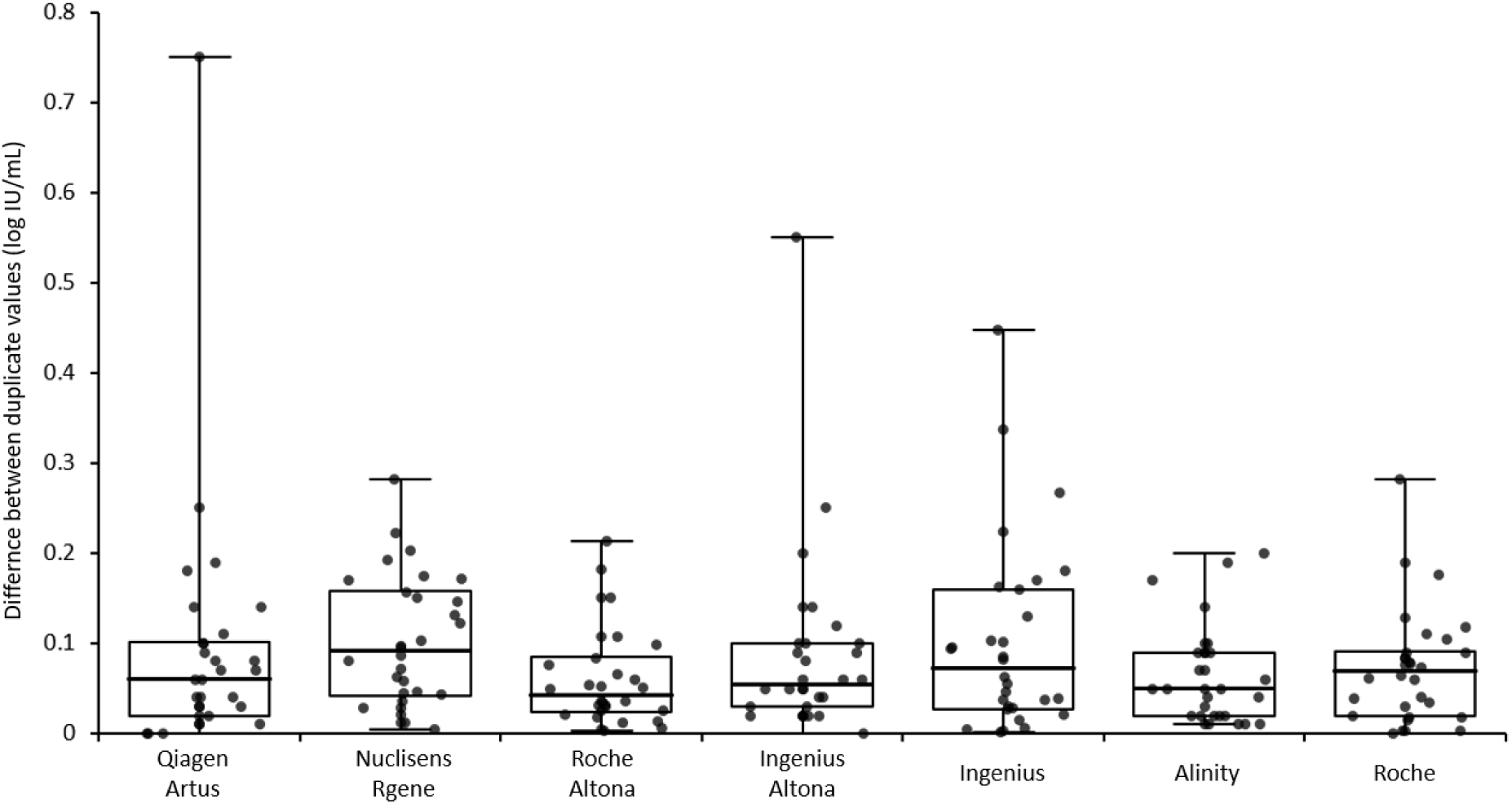
Box plot (median, IQR) of the difference between duplicate VL quantification for each system.

### Comparing the differences between the measured VLs and the means for the clinical specimens and the WHO/EDX standards

Each VL quantification obtained with each system individually for the clinical specimens and the WHO/EDX standards combined together were compared to each mean value obtained from all seven systems combined. Differences between the clinical specimens and the standards were then evaluated for each system (Figure 4). Significant differences between the clinical specimens and the standards VL measurement were observed for all the systems, except the Ingenius and Roche systems that appeared to quantify both the clinical specimens and the standard similarly (no statistically significant difference was observed). For the five remaining systems, the difference to the mean quantification was significantly different between clinical specimens and standards.

**Figure 4:**
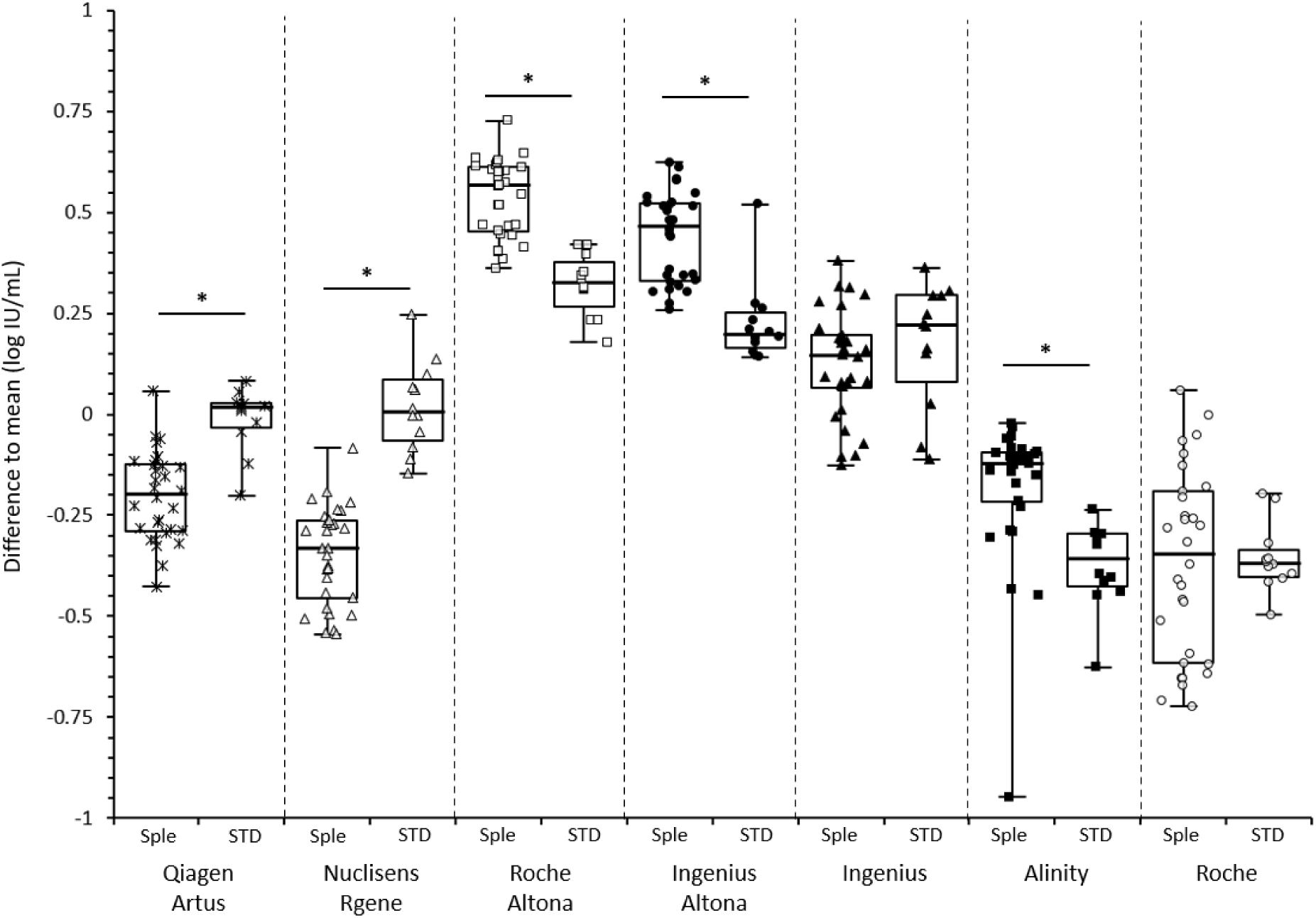
Difference to the mean VL for each system according to the type of samples: Clinical specimens (Sple) or WHO/EDX standards (STD). * indicates statistically significant differences.

### Quantification of the WHO and EDX reference standards

The Roche and Alinity systems appeared to be well calibrated with mean biases of 0.05 and −0.08 log_10_ IU/mL, respectively, between the nominal concentrations obtained and those of the WHO standards (Figure 5). In contrast, the Ingenius system showed the highest mean bias, over-quantifying the WHO standard by an average of 0.67 log_10_ IU/mL. For the remaining four systems the mean differences ranged from 0.32 to 0.650 log_10_ IU/mL. The Roche and Abbott systems accurately quantified the EDX standard with minor biases of −0.14 and −0.06 log_10_ IU/mL, respectively. However, the largest differences were observed with the Altona amplification-based systems with a mean bias reaching 0.62 for the Roche-Altona system.

**Figure 5:**
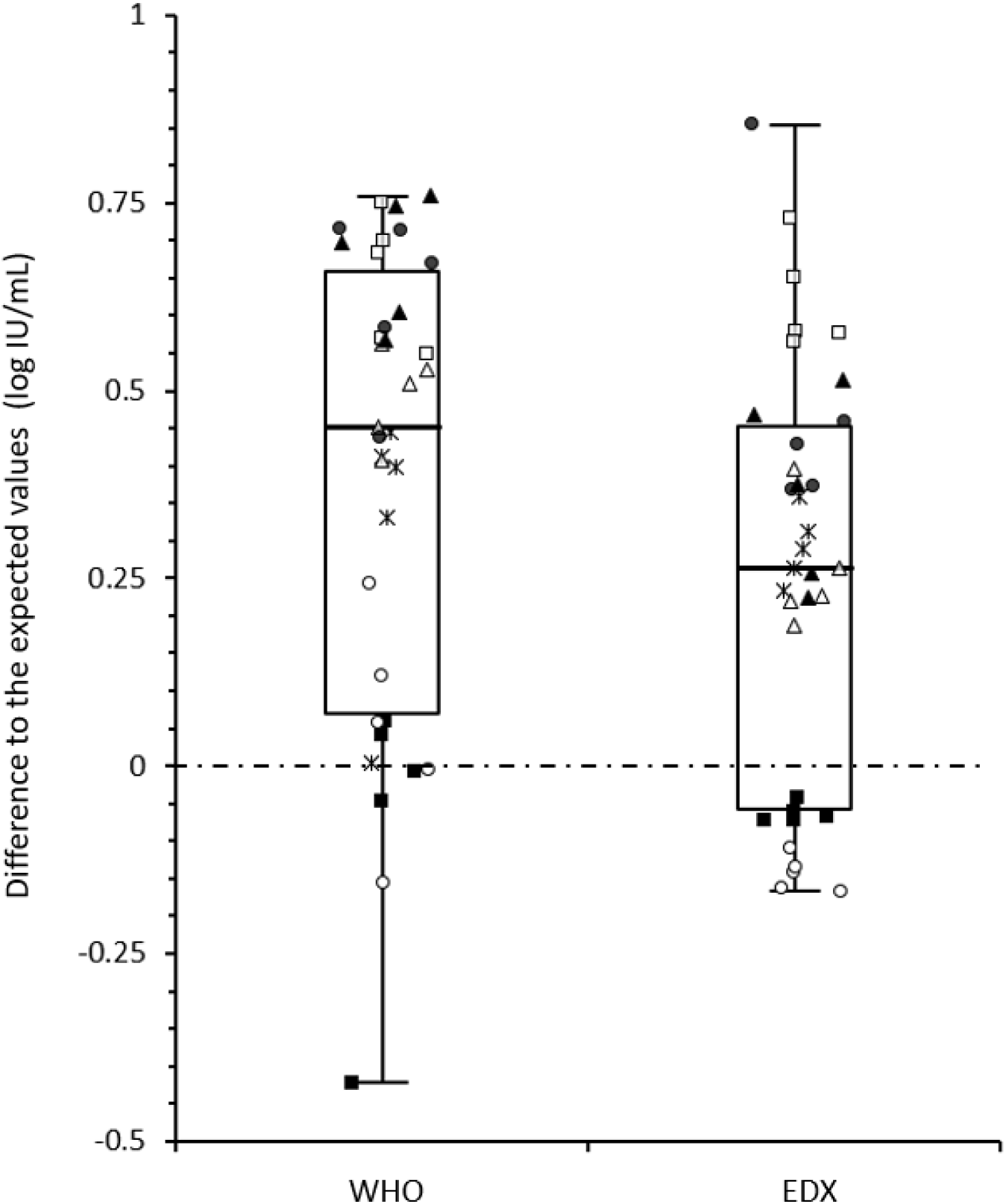
Difference in standard quantification by each system. Each symbol represents one standard concentration. Closed squares: Alinity; Closed triangles: Ingenius; Closed circles: Ingenius-Altona; Open triangles: Nuclisens-Rgene; Stars: Qiagen-Artus; Open circles: Roche; Open squares: Roche-Altona.

When all results were compared, a statistically significant difference (p=0.0217) was observed between the biases associated with the WHO standard and those obtained with the EDX standard. The biases for the EDX standard were generally lower than those for the WHO standard, regardless of the system.

### Benefit of recalibration with a universal reference standard

We investigated whether Ct values associated with standard materials (WHO or EDX) could be used to re-quantify the results initially obtained with each system’s in-kit calibration, with the goal of further standardizing VL measurements across systems (Figure 6). While the distribution of VL concentrations reached a median of 1.02 log_10_ IU/mL [IQR: 0.47] for all clinical specimens with the in-kit calibrators, it decreased after recalibration with both reference standards: to 0.86 log_10_ IU/mL [IQR: 0.21] (p<0.001) with WHO, and to 0.71 log_10_ IU/mL [IQR: 0.27] (p<0.001) with EDX.

**Figure 6:**
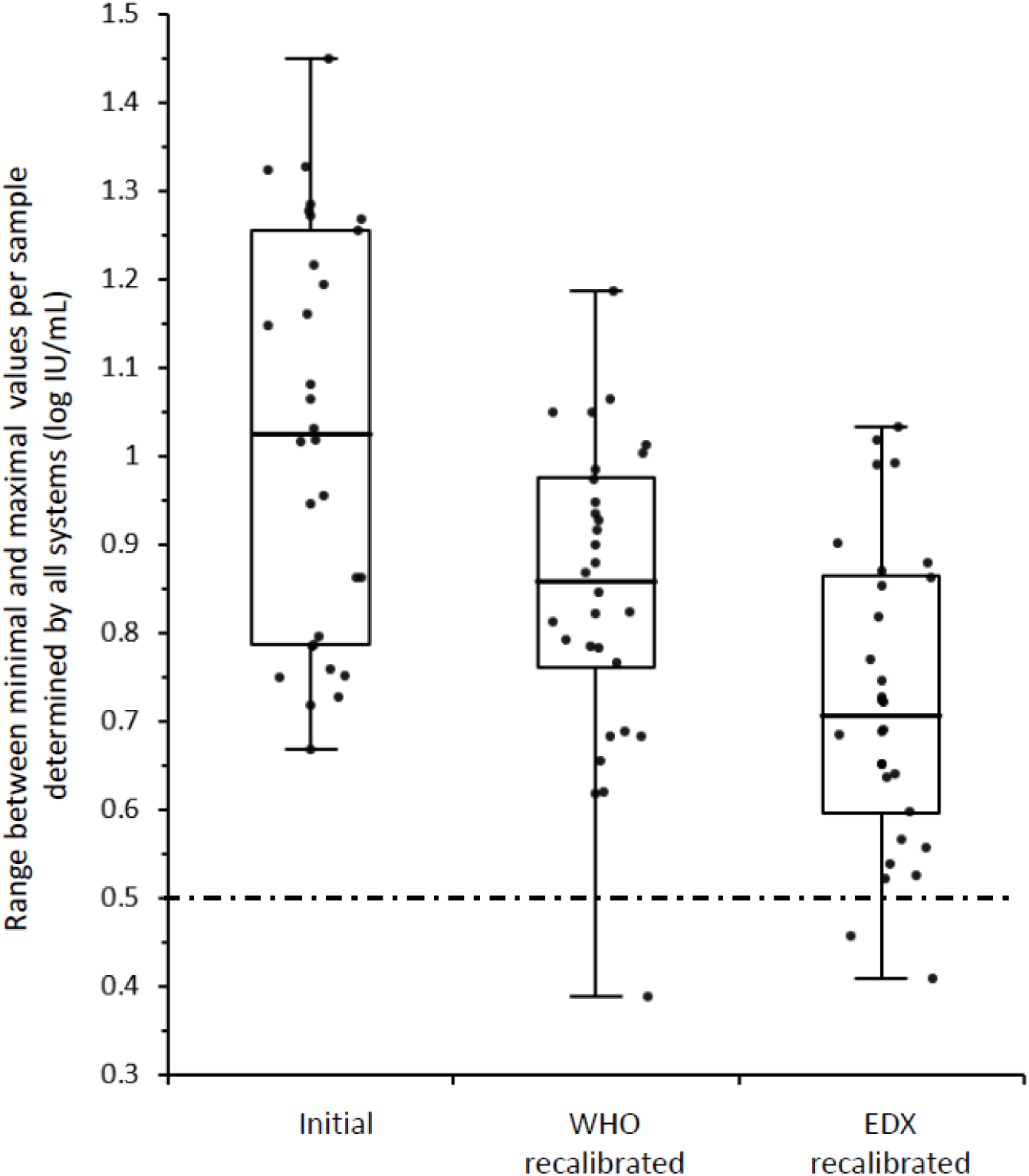
Scattering of clinical specimen VL measurements before and after recalibration with the WHO and EDX standards.

As expected, recalibration with either WHO or EDX standards also resulted in a reduction in the variation of VLs from the expected value for each standard (Table 1)

**Table 1:**
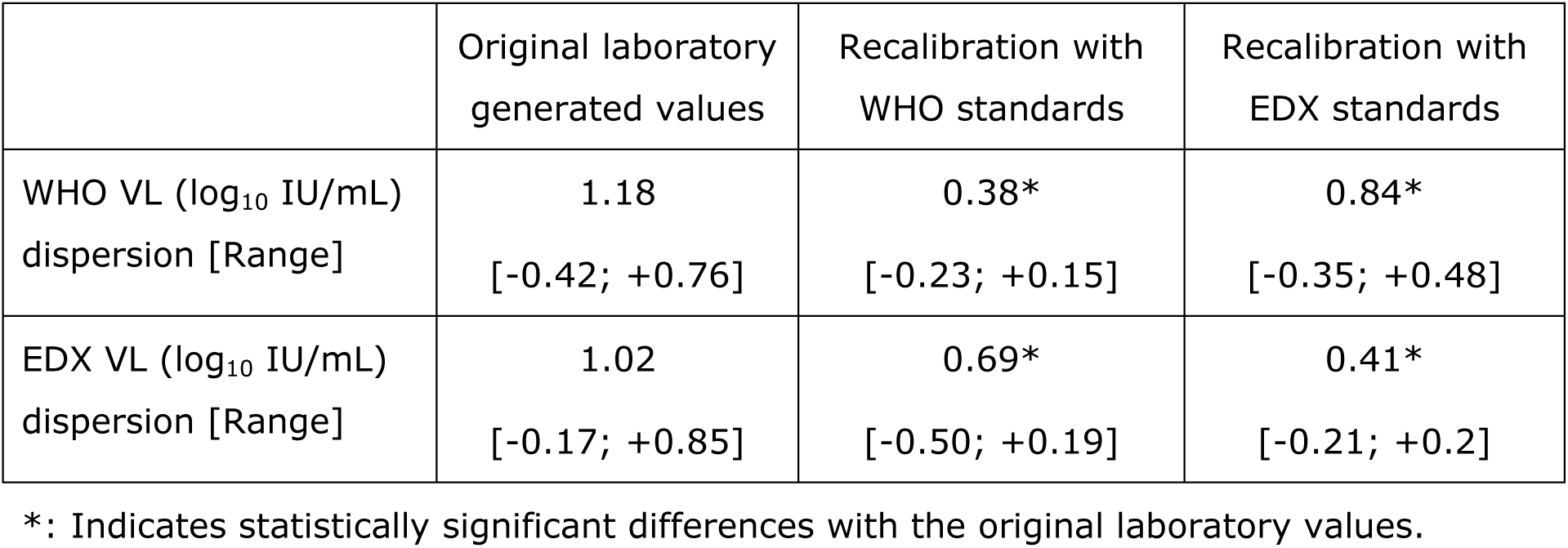
Inter-laboratory dispersion of standard CMV viral load values in the original set of results and after recalibration with WHO or EDX standards.

**Supplementary figure:**
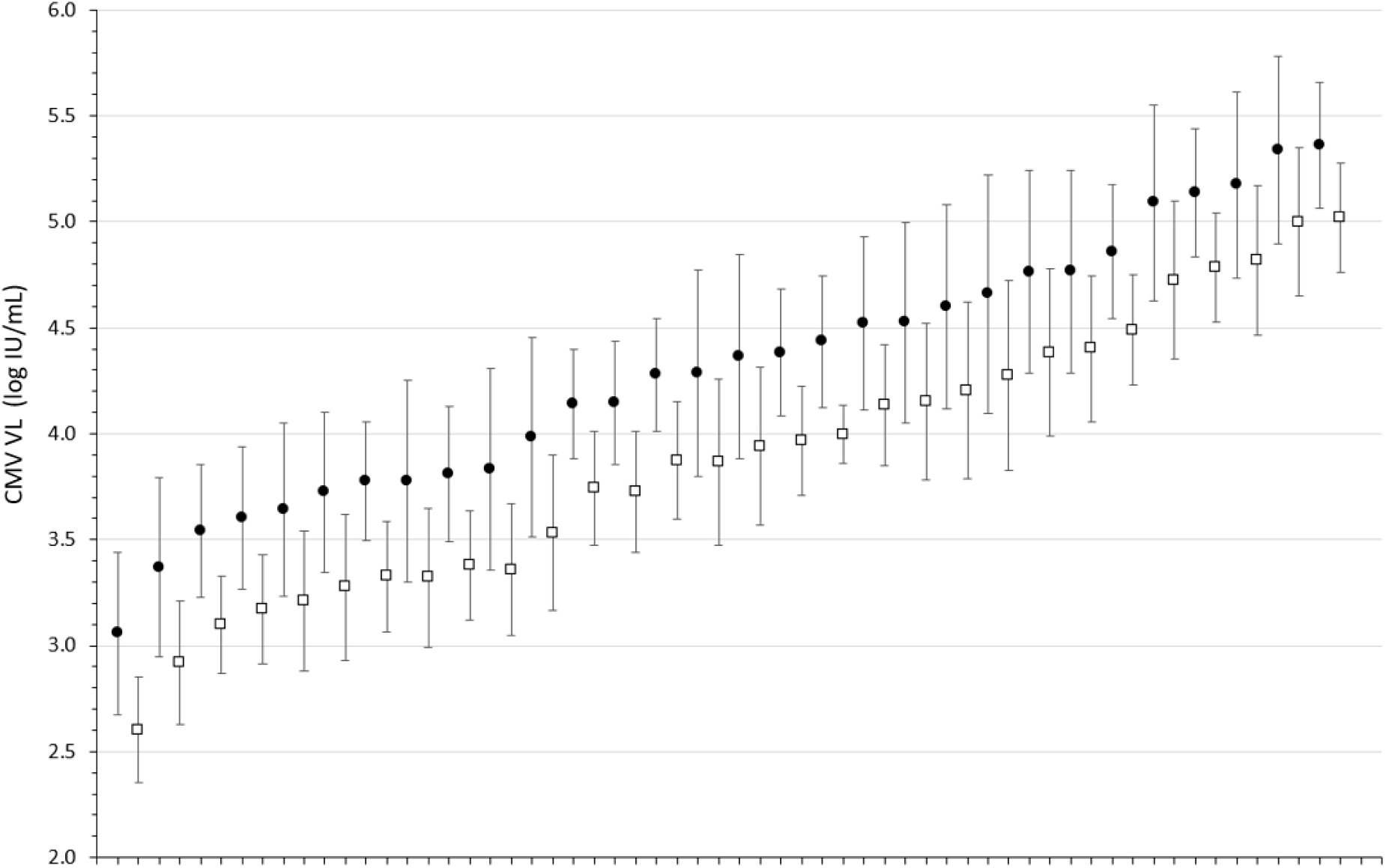
Mean VL measured by all the systems combined for each panel member. Error bars represent the standard deviation. Closed circle: In-kit calibration; Open square: After recalibration against the WHO standard.

## Discussion

The aim of this multicenter study was to evaluate progress towards harmonization of CMV VL quantification under routine testing conditions, taking into account the diversity of platforms and CMV assays used in France.. Of the thirteen commercial CMV assays on the market at the time of the study, the vast majority, if not all, used in-kit standards of plasmids or amplicons. Notably, this study represents the first documented use of the WHO international standard for recalibration alongside a commercial secondary standard (EDX). Our results, in agreement with recent literature, show that considerable efforts are still required to achieve full standardization across the different systems tested. We also propose further insights into the necessary steps to achieve a higher level of harmonization, similar to the standardization observed for other bloodborne viruses such as HIV and HBV (11).

Overall, all systems tested showed good performance with strong correlation between them, excellent linearity in quantification of the WHO standard and a commercial EDX standard (Bio-Rad), and good consistency in testing duplicates.. It is noteworthy that no false positives were observed despite the presence of highly viremic samples randomly mixed with negative samples, highlighting the overall improvement in laboratory sample handling and molecular biology expertise over the past decades.

Except for VLs near the lower limit of quantification reported by some of the systems, reproducibility, as measured by the within-duplicate quantification difference, was typically less than 0.20 log_10_ IU/mL (Figure 3). However, the IQR calculated for each clinical specimen quantified by all the systems used in the study varied from 0.44 to 1.05 log_10_ IU/mL, with quantification differences ranging from 0.67 to 1.45 log_10_ IU/mL (Figure 2), a variability consistent with findings by other authors (Peddu et al. 2020). In contrast, the IQR for the quantified WHO standard with the different systems was narrower, ranging from 0.07 to 0.43 log_10_ IU/mL, and the difference in quantification for each point ranged from 0.22 to 0.67 log_10_ IU/mL (Figure 1). Interestingly, the quantification variability was less pronounced for the standards (IQR=0.13 log_10_ IU/mL) than for the clinical specimens (IQR=0.47 log_10_ IU/mL; p=0.0142). With respect to the expected performance of the techniques against the WHO reference standard (Figure 5), the smallest mean differences from the expected values were observed with Abbott (−0.07 log_10_ IU/mL) and Roche (0.05 log_10_ IU/mL), while Roche-Altona (0.65 log_10_ IU/mL), Ingenius-Altona (0.63 log_10_ IU/mL) and Ingenius (0.68 log_10_ IU/mL) were the systems that provided the most discrepant results. These observations underscore that while improvements have been made, standardization of results is still lacking when calibrating against common standards, with the exception of integrated systems. To make progress toward robust quantification, manufacturers may need to design their in-kit standards appropriately. This may be particularly critical given that most of these in-kit standards are made of plasmids, and it has been reported that standards made of circular plasmids can lead to an overestimationof 1.3 log_10_ in quantitative PCR measurements (Hou et al. 2010). While accurate calibration may not solve the entire problem, it can at least reduce the variation that is clearly related to the specific system used.

Since we could not compare quantification to a gold standard, we chose to compare all quantifications to the mean VL obtained from the seven systems used in this study. Surprisingly, the differences observed were not the same for clinical specimens and the WHO/EDX standards. For all systems except Roche and Ingenius, the differences from the mean values for the clinical specimens and the WHO/EDX standards were statistically significant. More specifically, for two systems (Qiagen-Artus and Nuclisens-Rgene), the difference was greater for the WHO/EDX standards than for the clinical specimens while the opposite was true for the other systems. These results suggest that clinical specimens and WHO/EDX standards are not quantified in the same manner by all the systems. Our observation may be attributed to the different composition of these samples: the WHO/EDX standards essentially consist of viral culture supernatants which differ significantly from the highly heterogeneous CMV-positive plasma found in clinical specimens and which contains circulating fragments of degraded CMV genome along with a minority of intact virions (12–15). Several studies have shown that the heterogeneous nature of the CMV-positive plasma, characterized by fragmented circulating CMV DNA, results in highly variable VL measurements. This variability, as evidenced by external control assessment programs, highlights the inconsistencies introduced at various analytical stages, including extraction and amplification (10, 16, 17). In contrast, cell culture supernatants primarily contain intact virions, which are a more uniform material and certainly less complex than plasma. As a consequence, the differences in extraction procedures and amplicon lengths between the different systems have a greater impact on the clinical specimens, and the results were more accurate on the WHO/EDX standards than on the clinical specimens. Because the WHO standard does not fully reflect the clinical specimens, its design should be revised, as suggested by Hayden et al. (18).

We compared the results of all techniques on WHO and EDX standards. The EDX standard is a commercial secondary standard composed of virus culture supernatant diluted in CMV-free plasma and calibrated against the WHO standard. Although the assigned values of both the WHO and EDX standards were very similar, the mean bias was slightly lower with the EDX standard than with the WHO standard (0.26 vs 0.39, p=0.0217, Figure 5). There was no simple explanation for this observation, except perhaps, the number of culture passages which could potentially affect the fragmentation of the CMV genome.

Since the WHO and EDX standards had been included in the panel and quantified in a blinded manner using the in-kit standards, we could, in a second step, compare the raw results (Ct values) obtained from all the systems after recalibration with the WHO and the EDX standards. As shown in the Supplementary Figure and Figure 6, recalibration tended to slightly reduce the heterogeneity of VL measurements for each clinical sample and the standards.. However, even after recalibration, the VL variation for each sample remained mostly above 0.5 log_10_ IU/mL. Thus, a strategy consisting of using identical standards on different systems is unlikely to solve the problem of inter-laboratory commutability.

Inter-system variability in CMV DNA quantification has been documented for over a decade and remains the primary reason for the lack of commutability (9, 10, 16, 19). The observed discrepancies between the systems have been attributed to numerous factors, such as the nucleic acid extraction method, the amplification target, and the quantitative calibrator used to calculate the final results (20). With this in mind, we deliberately designed this study to include systems that reflect the technical diversity found in hospital laboratories. As shown in Supplementary Table 1, the input sample volume equivalent (ISVE) ranged from 40 to 202 across the systems, highlighting the heterogeneity of the entire process from the sample volume to the amplified material (21). ISVE is one parameter that can influence the results (14). Interestingly, the two integrated systems Abbott and Roche, that produced comparable results across the entire panel of samples, had similar ISVE (around 200) and were both the most accurately calibrated against the WHO standard.

Some limitations of this study should be acknowledged. First, unlike other clinical markers, there is no gold standard method to serve as an absolute reference. Therefore, we relied on the mean VL quantification from all systems for each sample, which carries a risk of bias if one method produces outlier values. Second, while many laboratories in France monitor CMV VL in whole blood samples, the panel in this study consisted of plasma samples, requiring the participating laboratories to adapt their procedures accordingly. Given the technical challenges, it was not possible to generate a consistent whole blood panel but no laboratories reported issues in processing the plasma specimens. Third, the panel was generated from frozen clinical samples and it was not possible to assess the influence of the freeze-thaw cycles on material quality. However, all participants worked with identical materials, limiting the risk of bias. Finally, ideally all participants should have been provided with identical recalibration sets, but not all systems allow calibration against external reference material. Therefore, we decided to perform the recalibration by calculating the new quantification values based on the WHO/EDX standards, which were initially quantified blindly. We felt that this was the most unbiased approach available under the circumstances. Several studies have shown that CMV DNA in clinical specimens does not primarily circulate as whole viruses but rather as small fragments, typically less than 300 base pairs in length (12–15, 22). Depending on the extraction process, some techniques may preferentially purify certain DNA fragment lengths over others, contributing to heterogeneity in VL quantification. This heterogeneity may not be uniform across the entire quantification range, and smaller fragments (often resulting from the lysis of infected cells) may be more common in clinical specimens with low VL. These variations in extraction efficiency based on DNA fragment size contribute to quantification bias between systems, a discrepancy further exacerbated by the fact that different systems target varying amplicon sizes and CMV DNA regions. In contrast, reference materials, such as those from the WHO or commercial sources, are derived from virus culture supernatants, which consist primarily of whole viruses with full-length CMV DNA. This homogeneity explains why systems using WHO/EDX reference materials show less bias than those using clinical specimens. In addition, new antiviral treatments, such as letermovir, may differentially affect CMV DNA quantification methods and should be considered when interpreting results (23).

In this study, we demonstrate that the use of a single source of calibration material somewhat reduces the inter-laboratory bias, but has a limited impact on the overall variability of results between individual clinical specimens. The current WHO international reference standard and commercially derived secondary counterparts may not be perfectly suited for such an application aimed at standardizing CMV VL quantification, as their composition does not accurately reflect the circulating CMV material found in the blood of infected patients. As demonstrated by Hayden et al. the use of a fragmented reference material may improve commutability between laboratories, and the development of such materials should be enabled (18). However, it is essential to first elucidate the fragmentation pattern of the CMV genome which may differ depending on the clinical situation.

This study used the commercial EDX CMV standard (Bio-Rad) as a secondary reference material, which is also evaluated in a study by Hayden et al. (24). The performance of the EDX standard closely matched that of the WHO IS, not only because EDX is calibrated against WHO but also because it is quantified by digital droplet PCR (Bio-Rad), a method that Hayden et al. suggest is important for improving harmonization in CMV quantification (24).

As emphasized in various national and international guidelines, the recommendation to monitor patients by consistent use of a single system should be prioritized, while inter-laboratory comparison of CMV VL is strongly discouraged (25). However, this recommendation is often impractical in clinical practice. Therefore, it would be valuable to explore the feasibility of using the same independent run controls across the different systems used to monitor a given patient’s CMV VL. This approach could help to mitigate and control bias between these systems, and potentially provide a means to quantify it as well.

In conclusion, we are still far from achieving perfect inter-laboratory commutability of CMV VL results. As emphasized in numerous guidelines over the years, it is still advisable to monitor a patient in a single laboratory or at least in different laboratories using the same assay (26, 27). Our findings suggest that integrated systems may help to reduce the risk of obtaining divergent results.

## Acknowledgments

Christophe Béna with INO Specimens BioBank (France)

## Author Contribution

- Conceptualization: VT
- Formal analysis: VT
- Investigation: DG, JBR, CR, VT
- Acquisition of data: DB, ASL, RG, BG, CA
- Methodology: DG, JBR, VT
- Resources: DG, JBR
- Supervision: VT
- Writing ± original draft: VT, DG, DB

---

**Supplementary table:** Characteristics of the different workflows used at each site.

| Sites | CH<br>Annecy (F) | CHU<br>Clermont Ferrand (F) | CHU Cochin<br>Paris (F) | CHU<br>Grenoble (F) | CHU Pitié-Salpêtrière<br>Paris (F) | CHU<br>Rennes (F) | Bio-Rad<br>Fort-Worth (USA) |
| --- | --- | --- | --- | --- | --- | --- | --- |
| Platform | Ingenius/Altona<br>RealStar 1.0 | Elitech Ingenius<br>CMV ELITE MGB | Abbott<br>Alinity m | BioMerieux<br>CMV R-GENE | QIAAsymphony<br>Qiagen CMV QS-RGQ | Flow Roche/Altona<br>RealStar 1.0 | Roche Cobas CMV |
| Integrated | No | Yes | Yes | No | No | No | Yes |
| **1) Extraction** |  |  |  |  |  |  |  |
| Name: | Elite Ingenius SP200<br>extraction reagent | ELITE InGenius | CMV AMP Kit | Nuclisens<br>(eMag) | QIAAsymphony<br>DNA mini Kit | MP96 Roche | Cobas 6800 |
| Starting vol. (µL): | 200 | 200 | 500 | 200 | 200 | 200 | 350 |
| Elution vol. (µL): | 50 | 100 | 60 | 50 | 90 | 100 | 50 |
| Concentration factor: | 4 | 2 | 8,3 | 4 | 2,2 | 2 | 7 |
| **2) Amplification** |  |  |  |  |  |  |  |
| Name : | Altona realstar<br>CMV PCR kit 1.0 | CMV ELITE MGB Kit<br>ELITE InGenius | CMV AMP Kit | CMV R-gene | artus CMV QS-RGQ | Altona RealStar<br>CMV PCR Kit 1.0 | Cobas 6800 |
| Extracted NA (µL): | 10 | 20 | 24,2 | 10 | 20 | 10 | 27 |
| ISVE* | 40 | 40 | 202 | 40 | 44 | 20 | 189 |
| LLD (IU): | 0.249 to 0.943<br>IU/µL (/PCR reaction) | 250 IU/ml | 30 IU/mL | 89 IU/mL | 70 IU/mL | 605 IU/mL | 34.5 IU/mL |
| Range: | 1.21 to 1.21E+09<br>IU/µL (/PCR reaction) | 178 to 10E+09<br>IU/ml (WB) | 30 to 10E+09<br>IU/ml | 385 -385 000<br>IU/mL | 130 - 1.6 x 10E+08<br>IU/mL | 1.21 to 1.21E+09<br>IU/µL (/PCR reaction) | 34.5 - 1E+07<br>IU/mL |
| Target gene: | ? | MIEA | UL34 & 80.5 | UL83 | MIE | ? | gB (UL55) |

\*Input sample volume equivalent (ISVE) = starting volume\*[template volume/elution volume]

## Bibliography

1. Griffiths P, Reeves M. 2021. Pathogenesis of human cytomegalovirus in the immunocompromised host. 12. Nat Rev Microbiol 19:759–773.

2. Razonable RR, Hayden RT. 2013. Clinical Utility of Viral Load in Management of Cytomegalovirus Infection after Solid Organ Transplantation. Clin Microbiol Rev 26:703–727.

3. Kotton CN, Kamar N. 2023. New Insights on CMV Management in Solid Organ Transplant Patients: Prevention, Treatment, and Management of Resistant/Refractory Disease. Infect Dis Ther 12:333–342.

4. Haidar G, Boeckh M, Singh N. 2020. Cytomegalovirus Infection in Solid Organ and Hematopoietic Cell Transplantation: State of the Evidence. J Infect Dis 221:S23– S31.

5. Limaye AP, Babu TM, Boeckh M. 2020. Progress and Challenges in the Prevention, Diagnosis, and Management of Cytomegalovirus Infection in Transplantation. Clin Microbiol Rev 34:e00043–19.

6. Lazzarotto T, Chiereghin A, Piralla A, Gibertoni D, Piccirilli G, Turello G, Campanini G, Gabrielli L, Costa C, Comai G, La Manna G, Biancone L, Rampino T, Gregorini M, Sidoti F, Bianco G, Mauro MV, Greco F, Cavallo R, Baldanti F, AMCLI-GLaIT. 2020. Kinetics of cytomegalovirus and Epstein-Barr virus DNA in whole blood and plasma of kidney transplant recipients: Implications on management strategies. PLoS One 15:e0238062.

7. Rzepka M, Depka D, Gospodarek-Komkowska E, Bogiel T. 2022. Whole Blood versus Plasma Samples-How Does the Type of Specimen Collected for Testing Affect the Monitoring of Cytomegalovirus Viremia? Pathogens 11:1384.

8. Pang XL, Fox JD, Fenton JM, Miller GG, Caliendo AM, Preiksaitisa JK. 2009. Inter-laboratory Comparison of Cytomegalovirus Viral Load Assays. American Journal of Transplantation 9:258–268.

9. Hayden RT, Caliendo AM. 2020. Persistent Challenges of Interassay Variability in Transplant Viral Load Testing. J Clin Microbiol 58:e00782–20.

10. Preiksaitis JK, Hayden RT, Tong Y, Pang XL, Fryer JF, Heath AB, Cook L, Petrich AK, Yu B, Caliendo AM. 2016. Are We There Yet? Impact of the First International Standard for Cytomegalovirus DNA on the Harmonization of Results Reported on Plasma Samples. Clin Infect Dis 63:583–589.

11. Wirden M, Larrouy L, Mahjoub N, Todesco E, Damond F, Delagreverie H, Akhavan S, Charpentier C, Chaix M-L, Descamps D, Calvez V, Marcelin A-G. 2017. Multicenter comparison of the new Cobas 6800 system with Cobas Ampliprep/Cobas TaqMan and Abbott RealTime for the quantification of HIV, HBV and HCV viral load. J Clin Virol 96:49–53.

12. Boom R, Sol CJA, Schuurman T, Van Breda A, Weel JFL, Beld M, Ten Berge IJM, Wertheim-Van Dillen PME, De Jong MD. 2002. Human cytomegalovirus DNA in plasma and serum specimens of renal transplant recipients is highly fragmented. J Clin Microbiol 40:4105–4113.

13. Peddu V, Bradley BT, Casto AM, Shree R, Colbert BG, Xie H, Santo TK, Huang M-L, Cheng EY, Konnick E, Salipante SJ, Jerome KR, Lockwood CM, Greninger AL. 2020. High-resolution profiling of human cytomegalovirus cell-free DNA in human plasma highlights its exceptionally fragmented nature. Sci Rep 10:3734.

14. Leuzinger K, Hirsch HH. 2023. Amplicon size and non-encapsidated DNA fragments define plasma cytomegalovirus DNA loads by automated nucleic acid testing platforms: A marker of viral cytopathology? J Med Virol 95:e29139.

15. Tong Y, Pang XL, Mabilangan C, Preiksaitis JK. 2017. Determination of the Biological Form of Human Cytomegalovirus DNA in the Plasma of Solid-Organ Transplant Recipients. J Infect Dis 215:1094–1101.

16. Naegele K, Lautenschlager I, Gosert R, Loginov R, Bir K, Helanterä I, Schaub S, Khanna N, Hirsch HH. 2018. Cytomegalovirus sequence variability, amplicon length, and DNase-sensitive non-encapsidated genomes are obstacles to standardization and commutability of plasma viral load results. J Clin Virol 104:39–47.

17. Hirsch HH, Lautenschlager I, Pinsky BA, Cardenoso L, Aslam S, Cobb B, Vilchez RA, Valsamakis A. 2013. An International Multicenter Performance Analysis of Cytomegalovirus Load Tests. Clinical Infectious Diseases 56:367–373.

18. Hayden RT, Tang L, Su Y, Cook L, Gu Z, Jerome KR, Boonyaratanakornkit J, Sam S, Pounds S, Caliendo AM. 2019. Impact of Fragmentation on Commutability of Epstein-Barr Virus and Cytomegalovirus Quantitative Standards. J Clin Microbiol 58:e00888–19.

19. Hayden RT, Sun Y, Tang L, Procop GW, Hillyard DR, Pinsky BA, Young SA, Caliendo AM. 2017. Progress in Quantitative Viral Load Testing: Variability and Impact of the WHO Quantitative International Standards. J Clin Microbiol 55:423–430.

20. Hayden RT, Yan X, Wick MT, Rodriguez AB, Xiong X, Ginocchio CC, Mitchell MJ, Caliendo AM, for the College of American Pathologists Microbiology Resource Committee. 2020. Factors Contributing to Variability of Quantitative Viral PCR Results in Proficiency Testing Samples: a Multivariate Analysis. Journal of Clinical Microbiology 50:337–345.

21. Naegele K, Weissbach FH, Leuzinger K, Gosert R, Bubendorf L, Hirsch HH. 2023. Impact of nucleic acid extraction procedures on human papillomavirus (HPV) detection and genotyping. Journal of Medical Virology 95:e28583.

22. Cook L, Starr K, Boonyaratanakornkit J, Hayden R, Sam SS, Caliendo AM. 2018. Does Size Matter? Comparison of Extraction Yields for Different-Sized DNA Fragments by Seven Different Routine and Four New Circulating Cell-Free Extraction Methods. J Clin Microbiol 56:e01061–18.

23. Giménez E, Gozalbo-Rovira R, Albert E, Piñana JL, Solano C, Navarro D. 2024. Letermovir use may impact on the Cytomegalovirus DNA fragmentation profile in plasma from allogeneic hematopoietic stem cell transplant recipients. Journal of Medical Virology 96:e29564.

24. Hayden RT, Su Y, Tang L, Zhu H, Gu Z, Glasgow HL, Sam SS, Caliendo AM. 2024. Accuracy of quantitative viral secondary standards: a re-examination. J Clin Microbiol 62:e0166923.

25. Razonable RR, Inoue N, Pinninti SG, Boppana SB, Lazzarotto T, Gabrielli L, Simonazzi G, Pellett PE, Schmid DS. 2020. Clinical Diagnostic Testing for Human Cytomegalovirus Infections. J Infect Dis 221:S74–S85.

26. Razonable RR, Humar A. 2019. Cytomegalovirus in solid organ transplant recipients-Guidelines of the American Society of Transplantation Infectious Diseases Community of Practice. Clin Transplant 33:e13512.

27. Ljungman P, de la Camara R, Robin C, Crocchiolo R, Einsele H, Hill JA, Hubacek P, Navarro D, Cordonnier C, Ward KN, 2017 European Conference on Infections in Leukaemia group. 2019. Guidelines for the management of cytomegalovirus infection in patients with haematological malignancies and after stem cell transplantation from the 2017 European Conference on Infections in Leukaemia (ECIL 7). Lancet Infect Dis 19:e260–e272.

